# Genetic variation in *Pan* species is shaped by demographic history and underlies lineage-specific functions

**DOI:** 10.1101/280016

**Authors:** Sojung Han, Aida M. Andrés, Tomas Marques-Bonet, Martin Kuhlwilm

**Affiliations:** Institut de Biologia Evolutiva (Consejo Superior de Investigaciones Científicas–Universitat Pompeu Fabra), Barcelona, Spain.; Department of Genetics, Max Planck Institute for Evolutionary Anthropology, Leipzig, Germany.; UCL Genetics Institute, Department of Genetics, Evolution and Environment, University College London, London, UK.; National Centre for Genomic Analysis–Centre for Genomic Regulation, Barcelona Institute of Science and Technology, Barcelona, Spain.; Institucio Catalana de Recerca i Estudis Avançats (ICREA), Barcelona, Spain.

## Abstract

Chimpanzees (*Pan troglodytes*) and bonobos (*Pan paniscus*) are the closest living relatives of humans, but they show distinct behavioral and physiological differences, particularly regarding female reproduction. Despite their recent rapid decline, the demographic histories of the two species have been different during the past one to two million years, likely having an impact on their genomic diversity. Here, we analyze the inferred functional consequences of genetic variation across 69 individuals, making use of the most complete dataset of genomic variation in the *Pan* clade to date. We test to which extent the demographic history influences the efficacy of purifying selection in these species. We find that small historical effective population sizes (*N_e_*) correlate not only with small genetic diversity, but also with more homozygous deleterious alleles, and an increased proportion of deleterious changes at low frequencies. Furthermore, we exploit the catalog of deleterious protein-coding changes on each lineage to investigate the putative genetic basis for phenotypic differences between chimpanzees and bonobos. We show that bonobo-specific non-synonymous changes are enriched in genes related to age at menarche in humans, suggesting that the prominent physiological differences in the female reproductive system between chimpanzees and bonobos might be explained, in part, by putatively adaptive changes on the bonobo lineage.

## Introduction

Chimpanzees (*Pan troglodytes*) and bonobos (*Pan paniscus*) are two closely related species with an estimated divergence time of 1 to 2 million years ago (Mya) and share the most recent common ancestor with humans 5 to 13 Mya (Fischer et al. 2011; Langergraber and Prüfer 2012; Prado-Martinez et al. 2013; De Manuel et al. 2016). The population history of bonobos is marked by multiple population bottlenecks and a small long-term effective population size (Ne). They are also known to have been geographically isolated in central Africa for a long time (nowadays Democratic Republic Congo) (Thompson 2003; Takemoto et al. 2017), likely increasing their genetic homogeneity. This is in stark contrast to the history of chimpanzees, which inhabited a much wider region across Africa (from Tanzania to Guinea). Most chimpanzee subspecies, except of the western population, are thought to have had a larger population size and to have maintained a greater level of genetic diversity compared to bonobos. For example, central chimpanzee *N_e_* is four times larger, and western chimpanzee *N_e_* is 1.4 times larger, than the *N_e_* of bonobos; these are the largest and smallest *N_e_* among chimpanzees, respectively (Kuhlwilm et al. 2016; De Manuel et al. 2016).

The two sister species have important differences in phenotypic traits and social dynamics. Chimpanzees show a clear hierarchy and frequent violence among males, while it has been suggested that bonobos show more egalitarian interactions (Nishida 1983; Goodall 1986; Furuichi 1987; Idani 1990), and female-female sexual interactions have been observed in bonobos but not chimpanzees (Kuroda 1980). Both bonobo and chimpanzee females display maximal or exaggerated sexual swellings in the perineal part around their ovulation period, which is a distinct feature of the *Pan* clade among the great apes (Nunn 1999). However, only bonobos have a prolonged maximal sexual swelling, where females appear to be estrous even when they are not ovulating (*e.g*. during lactating period, Thompson-Handler et al. 1984; Furuichi 1987). This has been suggested to be one reason why bonobos maintain peaceful social dynamics, by lowering the operational sex ratio (the temporal ratio of adult males to estrous females in the group, Randall Parish 1994; Furuichi 2011) and by strengthening female-female sexual interactions (as females are more attracted to other females that show sexual swellings, which builds female affiliative relationships and helps them have a strong social stance in their male-philopatric society, Ryu et al. 2015). Considering that the high social status of bonobo females is suggested to be a driving force of selection on non-aggressive males (Hare et al. 2012), the prolonged sexual swelling can be understood as a key aspect of the bonobo-specific nature. However, these traits in female reproduction have not yet been associated to genetic changes. On the physiological level, using urinary testosterone hormonal levels, it has been shown that female bonobos are on average three years younger than their chimpanzee counterparts when they experience the onset of puberty, which is an important clue for sexual development (Behringer et al. 2014). We propose that these physiological differences have a genetic origin, and identifying the genetic changes responsible for these traits would help us better understand the evolution of *Pan* species. Derived alleles in a lineage at high frequencies are the obvious candidate changes for such lineage-specific traits, and deleterious changes might be particularly relevant since they have an increased potential for functional impact.

On the other hand, whether different demographic histories influence the efficacy of purifying selection in populations has been a debated topic in the field of population genomics. Some studies have shown that in human populations with small *N_e_* and with population bottlenecks, genetic diversity is lower (number of heterozygous genotypes per individual; Lohmueller et al. 2008; Kidd et al. 2012; Torkamani et al. 2012; Hodgkinson et al. 2013) and the rate of random fixation of derived alleles is higher (proportion of fixed substitutions and number of homozygous derived alleles per individual; Lohmueller et al. 2008; Kidd et al. 2012; Torkamani et al. 2012) than in populations that have maintained a larger *N_e_*. These patterns are generally explained by the effects of genetic drift. However, for estimating the efficacy of purifying selection, it is crucial to consider the proportion of deleterious changes, particularly at high frequencies within the population. It has been inferred that almost half of all non-synonymous Single Nucleotide Polymorphisms (SNPs) in a single genome are deleterious (Subramanian and Lambert 2012). Such deleterious allele changes are, for example, predicted to alter protein function, which is typically strongly conserved across species, or are associated with an increased risk of disease; hence, they are proposed to be under purifying selection that reduces their probability of fixation in the population (Jukes and Kimura 1984). Slightly deleterious changes are particularly interesting because they are relatively well tolerated, can appear as polymorphisms (Henn et al. 2016), and thus could be informative about the effects of purifying selection. We expect a larger proportion of these slightly deleterious SNPs at high frequencies in a population with relaxed purifying selection pressure than in one with efficient purifying selection. This hypothesis has been tested in different species, including archaic and modern humans (Lohmueller et al. 2008; Castellano et al. 2014), dogs (Marsden et al. 2016), and rice (Liu et al. 2017).

Previous studies have analyzed the efficacy of purifying selection in the *Pan* clade: Using the exomes of central, eastern and western chimpanzees, (Bataillon et al. 2015) showed that the efficacy of natural selection correlates with past *N_e_*, in agreement with (Cagan et al. 2016) where lineage-specific selection signatures from all the great ape genomes led to the same conclusion. On the other hand, de Valles-Ibáñez et al. 2016 interpreted from the genomes of various great ape species, including eastern and Nigerian-Cameroon chimpanzees, that the load of loss-of-function (LoF) mutations, which probably have severe consequences, is not influenced by the demographic history. However, given the differences in demographic history across *Pan* subspecies, it is important to include all four known subspecies of chimpanzee, particularly the central and western chimpanzee populations, as well as bonobos.

In this study, we examine the genetic variation across 59 chimpanzee and 10 bonobo genomes (De Manuel et al. 2016) and infer the functional consequences of single-nucleotide changes. First, we analyze patterns of an accumulation of putatively deleterious derived alleles, with the goal of testing how their different demographic histories have shaped the distribution of such alleles. We stratified the alleles into heterozygous and homozygous genotypes per individual, and across functional categories, also in LoF mutations, which are the most disruptive type of deleterious changes. Second, we analyze changes that occur on the bonobo or the chimpanzee lineage. We prioritize these by using non-synonymous changes, and assess the enrichment in genes associated to different phenotypes in humans using the GWAS database. We interpret the enrichment in genes related to these traits as indication of a genetic basis for lineage-specific differences between the two *Pan* species.

## Materials and Methods

### Data Preparation

Chimpanzee and bonobo genomes used in this study were generated in previous studies (Prado-Martinez et al. 2013; De Manuel et al. 2016). The data consists of 59 chimpanzee and 10 bonobo genomes sequenced at high coverage, including four chimpanzee populations: 18 central, 20 eastern, 10 Nigerian-Cameroon and 11 western chimpanzees (Table S1). All genomes were mapped to the human genome assembly *hg19* (ENSEMBL GRCh37.75), and previously published genotype calls for single nucleotide changes (SNCs) – variable sites across all the genomes of the *Pan* populations we use in this study – on the autosomes were used (De Manuel et al. 2016). We applied the ENCODE 20mer uniqueness filter, and required all the heterozygous loci to be allele balanced, with ratios of the raw reads between 0.25 and 0.75, in order to exclude sites with potential biases or contamination. Across 69 individuals, the initial VCF file contained 36,299,697 loci in total. After applying all filters, we used 33,946,246 loci where at least one individual shows a reliable genotype, while only 5,795,261 loci have information for all 69 individuals. For the analysis with eight individuals per population at highest quality, we exclude samples which could have had a small percentage of human contamination. To do so, we counted positions where the human allele was only seen in one read in one individual, but a derived allele in at least five reads and all the other individuals, selecting individuals with less than 10,000 such positions across the genome. We then selected the individuals with the highest coverage from the remaining samples (Table S1). Sites without high-quality genotypes were not considered for these analyses. Furthermore, where stated, a subset of ten central chimpanzees for a fair comparison with bonobos was used (Table S1). In the analysis of GWAS genes described below, a more permissive population-wise filtering was applied, using sites at which at least 50% of individuals in each species (chimpanzee and bonobo) pass the above filters, and one of the lineages carries the derived allele at more than 90% frequency, while the other population carries less than 10% of the derived allele. Population-wise frequencies were calculated as proportions of the observed counts.

The functional effects for each SNC were inferred using the ENSEMBL Variant Effect Predictor (VEP) v83 (McLaren et al. 2016) on all segregating sites across the 69 individuals. Non-synonymous and synonymous SNCs were used as defined by VEP, and the following functional categories were retrieved from the ENSEMBL annotation: 5’ UTR, 3’ UTR, regulatory elements, splice sites, transcription factor binding sites, upstream variants, downstream variants. Genomic regions and positions were analyzed using *R/Bioconductor* (Huber et al. 2015) and the packages *biomaRt* (Durinck et al. 2005) and *GenomicRanges* (Lawrence et al. 2013). The catalog of variation and functional inference in *Pan* species is available under (will be available before publication).

### Terminology

“Derived change” refers to the allele state different from the reference allele state. In this study, the human genome was used as reference, and since it is an outgroup to the *Pan* clade it represents closely the ancestral state at *Pan*-specific mutations. “Ancestral state” refers to the allele state same as the human reference allele state. “Lineage-specific derived changes” refers to the derived changes fixed or almost fixed (>=90%) in one lineage (either chimpanzee or bonobo), where the other lineage appears fixed or almost fixed for the ancestral state (<10% derived). By doing this, we avoid the chances that we neglect the actual fixed sites which appear not fixed due to sequencing errors and include sites which are influenced by gene flow between populations.

### Deleteriousness assessment

We used the following methods to assess deleteriousness: Grantham score, C-score, GWAVA, SIFT, PolyPhen-2 and GERP score, to ensure our results are robust to the method used. The Grantham Score (Grantham 1974; Li et al. 1984) represents only the physical/chemical properties of amino acid changes. The C-score (Kircher et al. 2014) measures deleteriousness on a genome-wide scale for both coding and non-coding variants, integrating a variety of known functional information. GWAVA (Ritchie et al. 2014) predicts the functional impact of genetic variants, based on genome-wide properties. SIFT (Kumar et al. 2009) predicts the deleteriousness of amino acid changes, based on the degree of conservation inferred from sequence alignments. PolyPhen-2 (Polymorphism Phenotyping v2) (Adzhubei et al. 2010) predicts the possible impact of amino acid changes on the structure and function using both physical property and multiple sequence alignments. Finally, GERP (the Genomic Evolutionary Rate Profiling) scores (Davydov et al. 2010) compare, based on multiple alignments, the number of observed substitutions to the number of hypothetical substitutions given that they are neutral changes. They assume a deficit of observed substitutions as “Rejected Substitutions”, a natural measure of constraint on the element. For our analyses, we used precomputed base-wise scores for *hg19* (http://mendel.stanford.edu/SidowLab/downloads/gerp/). Neutral loci were defined as described in (Gronau et al. 2011).

### Analysis of non-synonymous lineage-specific SNCs

We used the software *FUNC* (Prüfer et al. 2007) to determine enrichment of Gene Ontology terms. We ranked the genes with the number of all lineage-specific non-synonymous changes, divided by the number of lineage-specific deleterious SNCs. In order to assess whether particular gene categories are enriched with lineage-specific deleterious SNCs in each lineage, we used the Wilcoxon rank test.

In order to formally assess an enrichment of lineage-specific non-synonymous changes in gene clusters associated with known phenotypic traits in humans, we retrieved genome-wide association data from the NHGRI-EBI GWAS Catalog (MacArthur et al. 2017), containing 2,385 associated traits. We analyzed all the associated genes in the data, which have protein-coding SNCs either on the chimpanzee or bonobo lineage, respectively. We used the permissive set of non-synonymous SNCs (Methods). For each trait from each study (“Disease trait”), we counted the number of non-synonymous SNCs on each lineage, and performed a significance test (G-test) against the total number of non-synonymous SNCs, compared to the total number of genes associated to the trait and the total number of protein-coding genes. We also performed a G-test between the total numbers of all non-synonymous SNCs on each lineage compared to the numbers of non-synonymous SNCs on each lineage falling in genes associated to the trait, to determine whether or not the difference between the two species is significant. In both cases, a P-value cutoff of 0.1 was applied. Finally, we performed an empirical enrichment test, by creating 500 random sets of genes with similar length as the genes associated to each trait (± 10% of the length of each gene), and counting the number of lineage-specific non-synonymous SNCs in each random set. Here, we require 90% of random sets to contain fewer non-synonymous SNCs in a given lineage than the real set of associated genes, to determine a trait significant. Finally, we filter for traits with at least 10 associated loci in order to restrict the analysis to multigenic traits, and we report only significant traits where at least three genes carry non-synonymous SNCs.

We screened the genes which have lineage-specific non-synonymous SNCs at high frequency, and confirmed their expression in chimpanzees and bonobos, using the dataset from (Brawand et al. 2011). The RNA sequencing data was mapped to the human genome assembly *hg19* using *tophat2* (Kim et al. 2013), and gene expression was estimated with *samtools* (Li et al. 2009) and *htseq-count* (Anders et al. 2015).

## Results

### Neutrality index in populations

We used inferred fixed derived (D) and polymorphic (P) non-synonymous (n) and synonymous (s) alleles in samples of equal size of eight individuals from each population to calculate a population-wise version of the Neutrality Index (NI) (Rand and Kann 1996), which is the ratio of *P_n_*/*P_s_* and *D_n_*/*D_s_*. The NI quantifies the direction and degree of natural selection, based on the assumption that synonymous substitutions are neutral. In the classical NI, a value greater than one indicates an excess of non-synonymous polymorphism over fixed derived sites, implying negative selection. A value lower than one might indicate an excess of fixed non-synonymous alleles and imply positive selection. We find that the NI across autosomes in western chimpanzees (1.51) is higher than in the other chimpanzee subspecies (1.19-1.28), caused by an excess of non-synonymous over synonymous polymorphisms (Fig. 1A, Table S2). This could be explained by reduced efficacy of purifying selection in this population as the result of a low *N_e_* (Eyre-Walker 2006). In bonobos, the NI is even higher than in non-western chimpanzees (1.36). The excess of potentially deleterious polymorphism in western chimpanzees is not exclusive for protein-coding changes, but exists also in different categories of non-coding sites in functional elements, such as 5’UTRs and the upstream and downstream regions of genes, compared to neutral changes across the genome (Fig. 1B, Table S2). This suggests that, also on non-coding loci, the efficacy of purifying selection is low in populations with small *N_e_* in the past.

**Figure 1.**
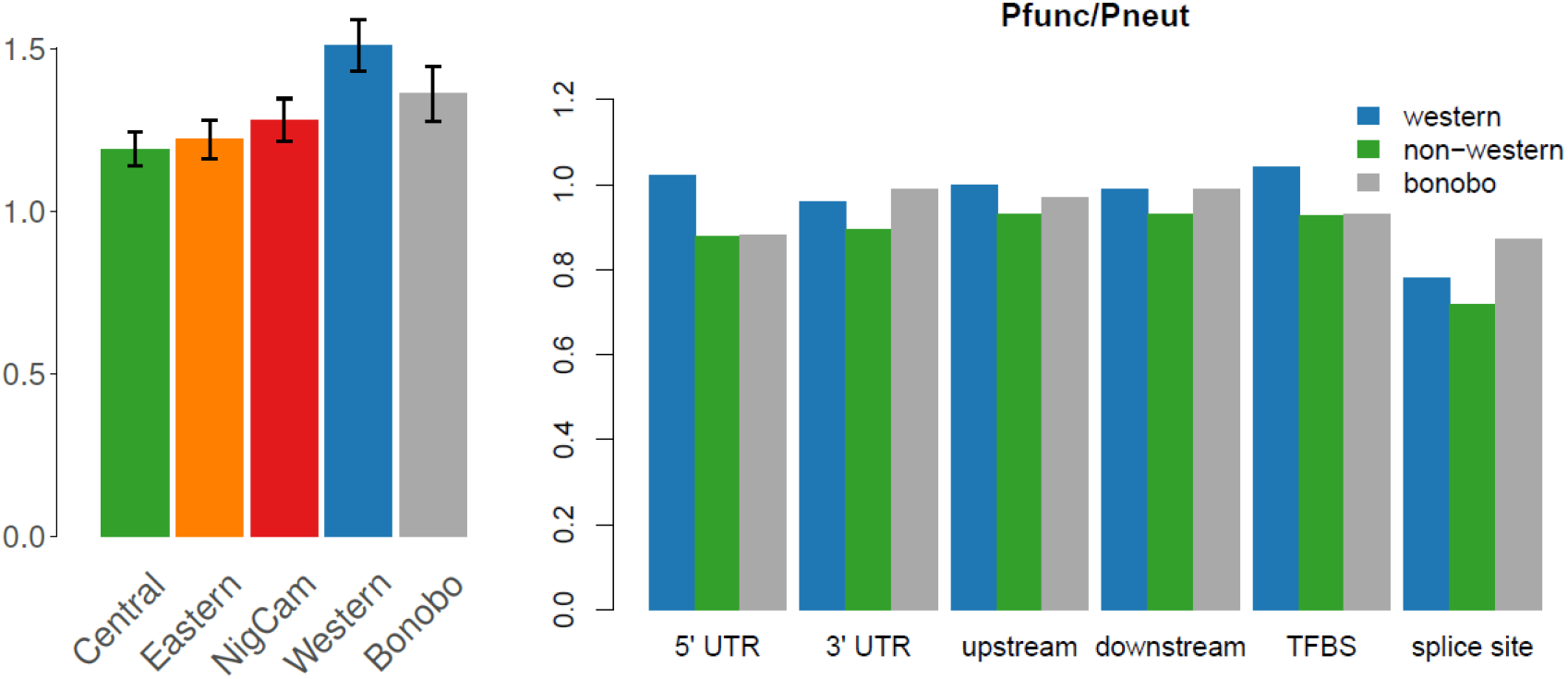
Patterns of functional and neutral alleles. A) Neutrality index in the five populations, averaged across autosomes. B) Genome-wide ratio of functional to neutral polymorphism (P_func_/P_neut_) in different functional categories, for western chimpanzees, bonobos and the average of three non-western chimpanzee populations.

### Distributions of deleterious changes

We analyzed the population-wise ratio of deleterious to neutral derived alleles across the site frequency spectrum (SFS) (Fig. 2), using eight individuals from each population. Deleterious alleles were defined each by GERP (Davydov et al. 2010) and PolyPhen-2 (Adzhubei et al. 2010), which represents genome-wide and protein-coding predictions, respectively. We generally observe a much higher proportion of deleterious derived alleles at high frequencies in bonobos than chimpanzees. When stratifying alleles into singletons vs. non-singletons, the deleterious-to-neutral ratio in non-singletons is highest in bonobos (0.96) than in chimpanzee populations (0.90-0.93). This suggests that bonobos, which experienced a long-term small *N_e_* and multiple bottlenecks since the split from chimpanzees (Prado-Martinez et al. 2013; De Manuel et al. 2016), have accumulated proportionately more deleterious alleles at high frequencies than any chimpanzee population. We interpret this as the long-term effect of low efficacy of purifying selection. At low frequencies, we observe all non-central chimpanzee populations having higher proportions of deleterious derived alleles, with western chimpanzee being particularly high. These patterns are generally similar using other deleteriousness estimates (Fig. S1).

**Figure 2.**
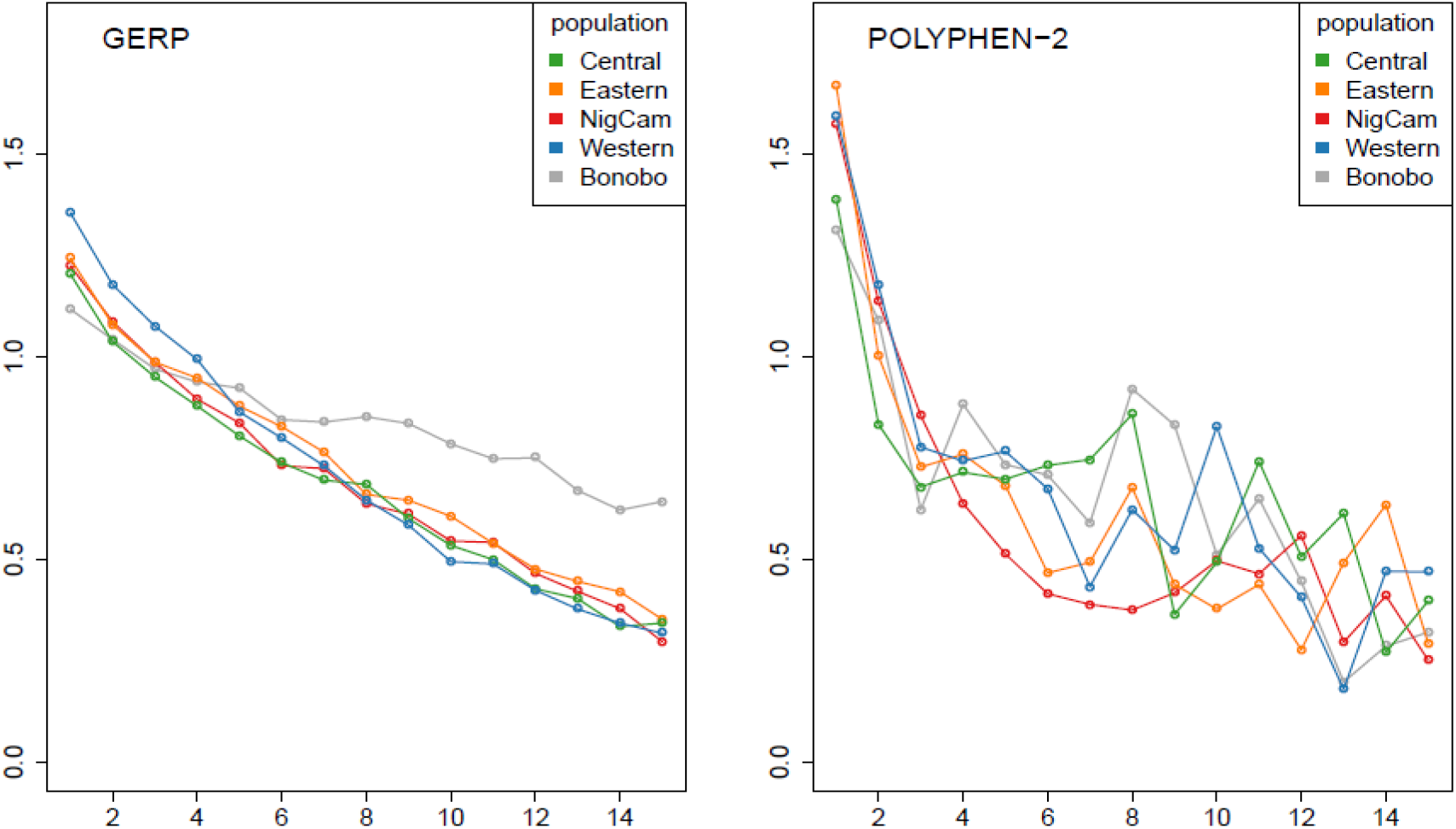
Ratio of Site-Frequency-Spectra (SFS) of deleterious to neutral derived alleles. Relative proportion of deleterious derived allele frequencies in each population, defined by GERP (Davydov et al., 2010) and POLYPHEN-2 (Adzhubei et al. 2010), respectively, compared to derived allele frequencies in neutral genomic regions, defined by (Gronau et al. 2011). For a fair comparison, 8 individuals from each population were chosen for the analysis.

Another way to assess the efficacy of purifying selection is to investigate the effects of population size on the individual mutational load. We estimated the mutational load, defined as the number of sites with putatively deleterious derived alleles per individual. We stratified them into two different classes, by counting either only heterozygous sites or only homozygous sites. When only heterozygous sites are considered, the western chimpanzee population shows the lowest level of mutational load among chimpanzee populations (Fig. 3A). This is significantly lower than in the other chimpanzee populations (P < 0.001; Wilcoxon rank test), and is largely due to their low genetic diversity. However, when only homozygous sites are considered, the mutational load of western chimpanzees increases drastically (Fig. 3B), and becomes significantly higher than that of central chimpanzees (P < 0.001; Wilcoxon rank test). This is probably because slightly deleterious alleles can more often reach high frequencies and be observed in homozygosity in populations with small *N_e_*, since purifying selection is less efficient. Regarding the other chimpanzee populations, the mutational load shows a gradient with a non-significant correlation trend with their long-term *N_e_* (De Manuel et al. 2016), negative in homozygous sites (r=−0.85 with p=0.07, Spearman correlation test) and positive in heterozygous sites (r=0.75 with p=0.15). This pattern is broadly similar throughout other classes of sites, including synonymous changes, in agreement with previous observations (Lohmueller et al. 2008) (Figs. S8-S13). This pattern appears less clear using C-score and GERP (r=−0.263 with p=0.738 and r=0.136 with p=0.864 respectively, Spearman correlation test). This might be due to the reference chimpanzee genome used in these methods being a western chimpanzee, as that could lower these scores for the derived changes in western chimpanzees. Still, the distributions of the heterozygous and homozygous mutational load of central chimpanzees are different from the other three chimpanzee populations in C-score and GERP (P < 0.0001 and P = 0.009 respectively; Wilcoxon rank test).

**Figure 3.**
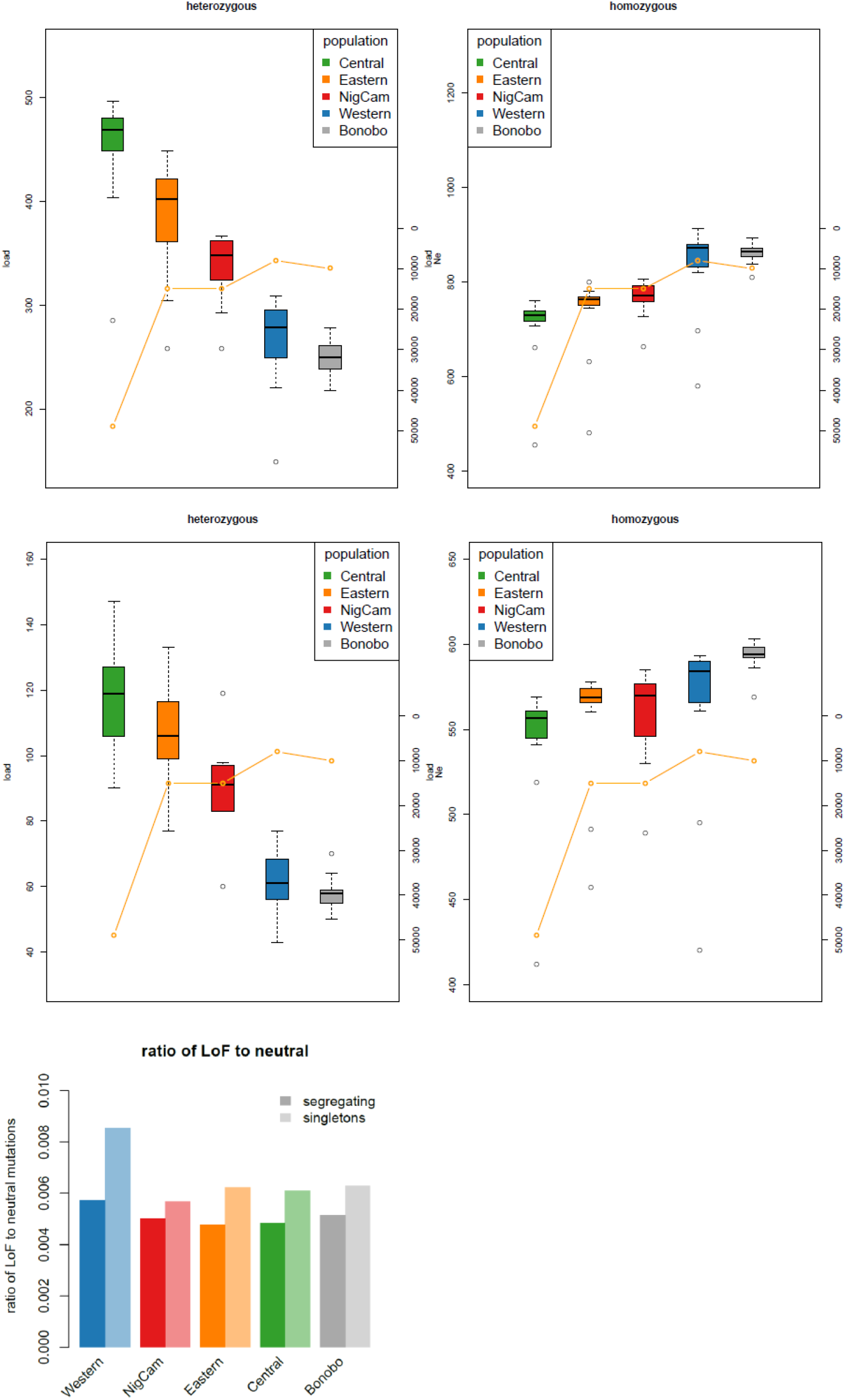
Mutational load and loss-of-function (LoF) variants. A) Individual mutational load in each population, considering only heterozygous sites (left). Mutational load considering only homozygous sites (right). These are the counts of deleterious derived alleles in each individual in each population, as defined by SIFT. B) Individual number of LoF variants, considering only heterozygous sites (left). Individual number of LoF variants, considering only homozygous sites (right). C) Ratio of LoF variants to neutral variants in each population.

### Protein-truncating variants

We also assessed the patterns in protein-truncating variants in each population by estimating the mutational load in LoF mutations, defined as the number of LoF derived alleles per individual. These loss-of-function (LoF) SNCs may be considered as most likely disruptive for protein function and hence a straightforward measure for evaluating the efficacy of purifying selection. In our data, the patterns of mutational load in LoF mutations follow the patterns in other categories of deleterious mutations (Fig. 3B). With higher *N_e_* the load tends to increase in heterozygous sites, although the correlation is not significant (r=0.786 with p=0.115). On average, chimpanzee individuals carry between 61 and 118 heterozygous LoF alleles, and between 547 and 560 homozygotes. In comparison, the average mutational load of LoF mutations in modern human populations is 85 for heterozygous sites (Lek et al. 2016), which is in agreement with previous observations of a slightly higher number of heterozygous LoF mutations in chimpanzees than humans, but smaller than in gorillas (Xue et al. 2015; de Valles-Ibáñez et al. 2016). We do not directly compare the number of homozygous sites to those in modern humans because in (Lek et al. 2016) only polymorphisms within the human lineage were used, while in our analysis, not only polymorphisms but also fixed differences between the two *Pan* species were measured (De Manuel et al. 2016).

When considering the same numbers of individuals per population, we observe twice as many fixed LoF mutations in western chimpanzees, and 2.7-fold as many fixed LoF mutations in bonobos, than in central chimpanzees. Conversely, the number of LoF singletons is three times higher in central chimpanzees than western chimpanzees, while the other chimpanzee populations are similar to the western chimpanzees, as expected by their background diversity. However, in western chimpanzees, the proportion of LoF mutations to neutral mutations is higher in polymorphisms than in fixed variants (P < 0.01; G-test). Also, the proportion of polymorphic LoF to neutral sites is higher in western chimpanzees than central chimpanzees (P = 0.012, G-Test). This effect is particularly pronounced in singletons (Fig. 3C), suggesting that LoF mutations are more often tolerated in western chimpanzees compared to the other populations, which could be due to less efficient purifying selection.

### An overview of lineage-specific changes

We assessed lineage-specific single nucleotide changes (SNCs) for their predicted functional effect, since these are candidates for functional changes explaining lineage-specific traits. In Table 1, we provide an overview of these SNCs, stratified by annotation category, when using ten individuals each from the bonobo and central chimpanzee population. This shows that bonobos have, on average, about two-fold more lineage-specific changes than central chimpanzees. Not surprisingly this number is three-fold higher when using 10 bonobos and the 59 chimpanzees (Table S3).

**Table 1.**
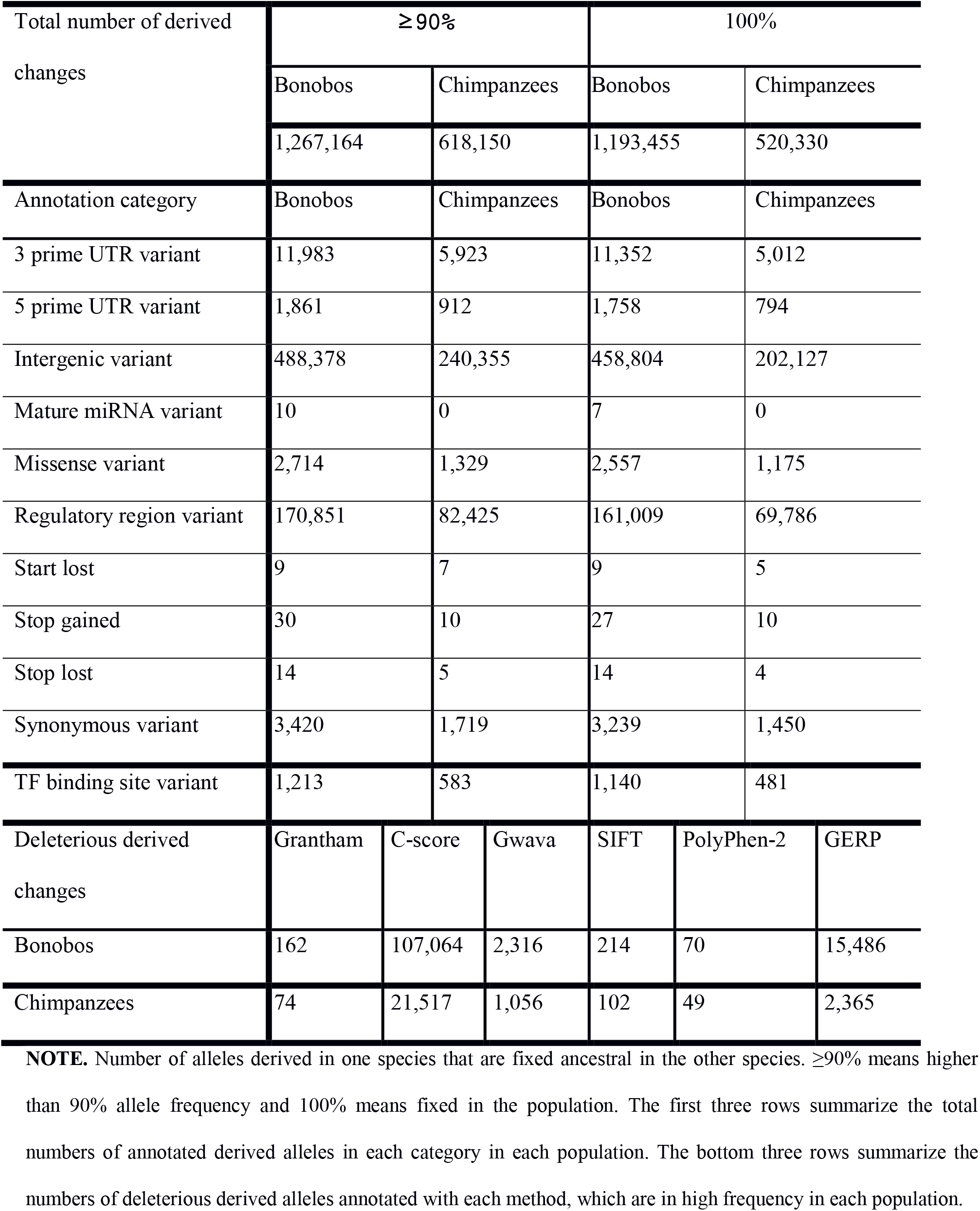
Summary statistics of the total number of lineage-specific changes in each species.

We assessed the deleteriousness of the lineage-specific SNCs as a proxy for functional and phenotypic consequences, using six different deleteriousness measures: Grantham score, C-score, GWAVA, SIFT, PolyPhen-2 and GERP score (Methods). We present a catalog of genes with lineage-specific deleterious SNCs among protein-coding changes, and of genes associated with 50 most deleterious lineage-specific SNCs in genome-wide inferences across the measures (Table S4, S5), in total affecting 78 genes in chimpanzees and 244 genes in bonobos. In bonobos, five of these genes are, according to the literature, involved in female reproduction: *ABCA13* (Nymoen et al. 2015), *ESPL1* (Gurvits et al. 2017), *KIF14* (Singel et al. 2014), *LVRN* (Nystad et al. 2014) and *MAP4* (Daisuke et al. 2009), while six are involved in male reproduction: *ACSBG2* (Fraisl et al. 2006), *GALNTL5* (Takasaki et al. 2014), *NME8* (Sadek et al. 2001), *WBP2NL* (Wu et al. 2007), *WDR63* (Hozumi et al. 2008) and *ZFHX3* (Hering et al. 2014). In chimpanzees, we identified only one gene related to female reproduction (*TOP2A*, Tubbs et al. 2009; Konecny et al. 2010) and one gene involved in male reproduction (*FANCL*, Wong 2003). Except for *LVRN* and *FANCL*, these genes are expressed in both *Pan* lineages. We also found that, in both species, multiples genes carry lineage-specific deleterious SNCs related to immunity, optic, heart and nervous system (Table 2).

**Table 2.** Bonobo-specific non-synonymous changes in menarche-related genes.

| **Position** | **Gene** | **Mutation** |
| --- | --- | --- |
| 1:14105326 | PRDM2 | T/A |
| 1:154574654 | ADAR | A/V |
| 1:177930850 | SEC16B | S/F |
| 1:66067125 | LEPR | S/A |
| 1:98348908 | DPYD | R/L |
| 2:137639763 | THSD7B | S/R |
| 2:141027837 | LRP1B | D/E |
| 2:141113964 | LRP1B | F/S |
| 2:141130669 | LRP1B | R/Q |
| 2:141215086 | LRP1B | S/T |
| 2:206911299 | INO80D | E/D |
| 2:206921468 | INO80D | L/V |
| 2:32631593 | BIRC6 | S/T |
| 2:44942458 | CAMKMT | A/T |
| 2:48027770 | MSH6 | K/R |
| 3:184049147 | EIF4G1 | E/K |
| 3:44841903 | KIF15 | T/I |
| 3:88205648 | C3orf38 | T/A |
| 4:104066209 | CENPE | K/E |
| 4:104070295 | CENPE | I/V |
| 4:104082402 | CENPE | L/V |
| 4:104098148 | CENPE | N/S |
| 4:106156559 | TET2 | R/K |
| 4:106156645 | TET2 | P/S |
| 4:106196843 | TET2 | T/P |
| 4:95129589 | SMARCA<br>D1 | N/S |
| 5:137713436 | KDM3B | L/I |
| 5:137756422 | KDM3B | N/S |
| 6:101296499 | ASCC3 | H/L |
| 6:128812928 | PTPRK | H/R |
| 6:148846475 | SASH1 | V/I |
| 6:170628088 | FAM120B | D/G |
| 6:84315463 | SNAP91 | A/V |
| 7:27867388 | TAX1BP1 | P/H |
| 8:1624699 | DLGAP2 | V/M |
| 8:2820076 | CSMD1 | Q/H |
| 8:35616858 | UNC5D | R/S |
| 8:35648008 | UNC5D | T/M |
| 8:53073977 | ST18 | I/V |
| 8:87423991 | WWP1 | N/D |
| 9:111798556 | TMEM245 | I/V |
| 9:120475206 | TLR4 | G/E |
| 9:6984389 | KDM4C | H/Y |
| 9:7170033 | KDM4C | T/I |
| 10:102684190 | FAM178A | R/C |
| 10:102684290 | FAM178A | C/Y |
| 10:13541765 | BEND7 | N/S |
| 11:122829992 | C11orf63 | P/S |
| 11:8474423 | STK33 | Q/E |
| 12:108638308 | WSCD2 | P/Q |
| 12:48073286 | RPAP3 | P/R |
| 13:42763236 | DGKH | A/V |
| 14:100847730 | WDR25 | V/I |
| 14:100847877 | WDR25 | R/C |
| 14:101349312 | RTL1 | E/G |
| 14:60950485 | C14orf39 | E/K |
| 14:94088581 | UNC79 | P/A |
| 14:94107526 | UNC79 | I/V |
| 15:48056211 | SEMA6D | I/M |
| 16:20380984 | PDILT | E/K |
| 16:72832557 | ZFHX3 | G/R |
| 17:2202857 | SMG6 | P/L |
| 17:2203679 | SMG6 | R/K |
| 17:46309238 | SKAP1 | V/I |
| 17:50212276 | CA10 | E/D |
| 20:37263383 | ARHGAP4<br>0 | G/S |
| 20:37275640 | ARHGAP4<br>0 | K/R |
| 20:43532655 | YWHAB | I/V |
| 20:43532673 | YWHAB | P/A |
| 20:62859360 | MYT1 | A/V |
| 22:50014212 | C22orf34 | G/R |
| X:109310617 | TMEM164 | V/I |
| X:8993603 | FAM9B | L/F |

Protein-truncating variants might have even stronger phenotypic effects. We find nine such SNCs in bonobos: one stop loss in *ACSM4*, and stop gains in *C12orf75*, the olfactory receptor *OR4N2, CABLES1, SIGLEC7, SEMA5B, B3GALNT1*, the breast cancer antigen *CEP85L* (Scanlan et al. 2001) and *TAAR2* (Table S6). On the other hand, chimpanzees carry only two LoF changes (in *ATP5L2* and *HEPACAM2*). To explore the putative effect of lineage-specific SNCs on phenotypes, we explored the NHGRI-EBI GWAS Catalog (MacArthur et al. 2017). We find one lineage-specific derived SNC that is almost fixed in bonobos but absent in chimpanzees, which is polymorphic in humans and associated with the trait “economic and social preference”. Chimpanzees carry the ancestral “risk allele” rs12606301-G. On the other hand, alleles that are at high frequency in chimpanzees, but absent in bonobos, are the human protective alleles rs17356907-G for breast cancer and rs3757247-A for both Vitiligo and type I diabetes, and the risk alleles rs1872992-G for “BMI-adjusted waist-hip ratio and physical activity measurement” and rs60945108-A for “physical activity”.

### Enrichment of deleterious and non-synonymous SNCs

To test if these lineage-specific deleterious SNCs are enriched in any particular gene family or pathway, we performed a formal Gene Ontology enrichment test. The results suggest that, on the bonobo lineage, among others, there is an enrichment in biological categories like “homophilic cell adhesion via plasma membrane adhesion molecules”, “steroid hormone mediated signaling pathway”, and “cell morphogenesis” (Table S10). On the other hand, in the chimpanzee lineage, we find an enrichment in categories such as “ionotropic glutamate receptor signaling pathway”, “positive regulation of GTPase activity”, and neuron-related categories like “positive regulation of axonogenesis” (Table S10).

In order to explore in further detail genes that are associated with polygenic traits, we performed a systematic enrichment screen for 2,385 traits from genome-wide association studies in humans (MacArthur et al. 2017), using a more permissive set of derived non-synonymous SNCs at high frequency (Methods). This analysis is based on the nearest genes to the associated site in humans, since most associated human SNPs are not segregating in the *Pan* dataset. We find 17 unique traits enriched for non-synonymous SNCs on the chimpanzee lineage, and five unique traits on the bonobo lineage (Table S11), among them “Menarche (age at onset)”. This suggests that in bonobos there is an enrichment of non-synonymous changes in genes affecting this female reproduction-related trait. Furthermore, we find an enrichment of such SNCs in genes associated to “Cognitive performance”, “Parkinson’s disease”, “Urinary albumin-to-creatinine ratio” and “Obesity-related traits”, which might reflect changes in bonobos related to cognitive abilities and metabolism. Genes with non-synonymous SNCs on the chimpanzee lineage are enriched, among others, for associations to traits involving body mass index and height, as well as “Schizophrenia” and “Loneliness”.

The finding of an excess of lineage-specific non-synonymous SNCs in genes associated with age at menarche suggests that we can identify genetic changes that may underlie the physiological differences between the two *Pan* species in terms of some female reproduction traits. We further investigated this trait using the 307 protein-coding genes associated to age at menarche in the most recent, most comprehensive meta-analysis of this trait to date (Day et al. 2017). Here, we observe a significant proportion of menarche-associated genes with bonobo-specific non-synonymous SNCs (P = 0.0025, G-Test, Table S12). This observation is even more pronounced when considering that 73 SNCs (Table 2) fall into these 55 candidate genes (P = 0.001, G-Test). This number of non-synonymous changes was not observed across 1,000 random sets of genes of similar length as the menarche-associated genes (Fig. 4). No enrichment of such changes is found on the chimpanzee lineage (P = 0.48, G-Test). We conclude that menarche-related genes seem to have acquired more non-synonymous SNCs than expected on the bonobo lineage.

**Figure 4.**
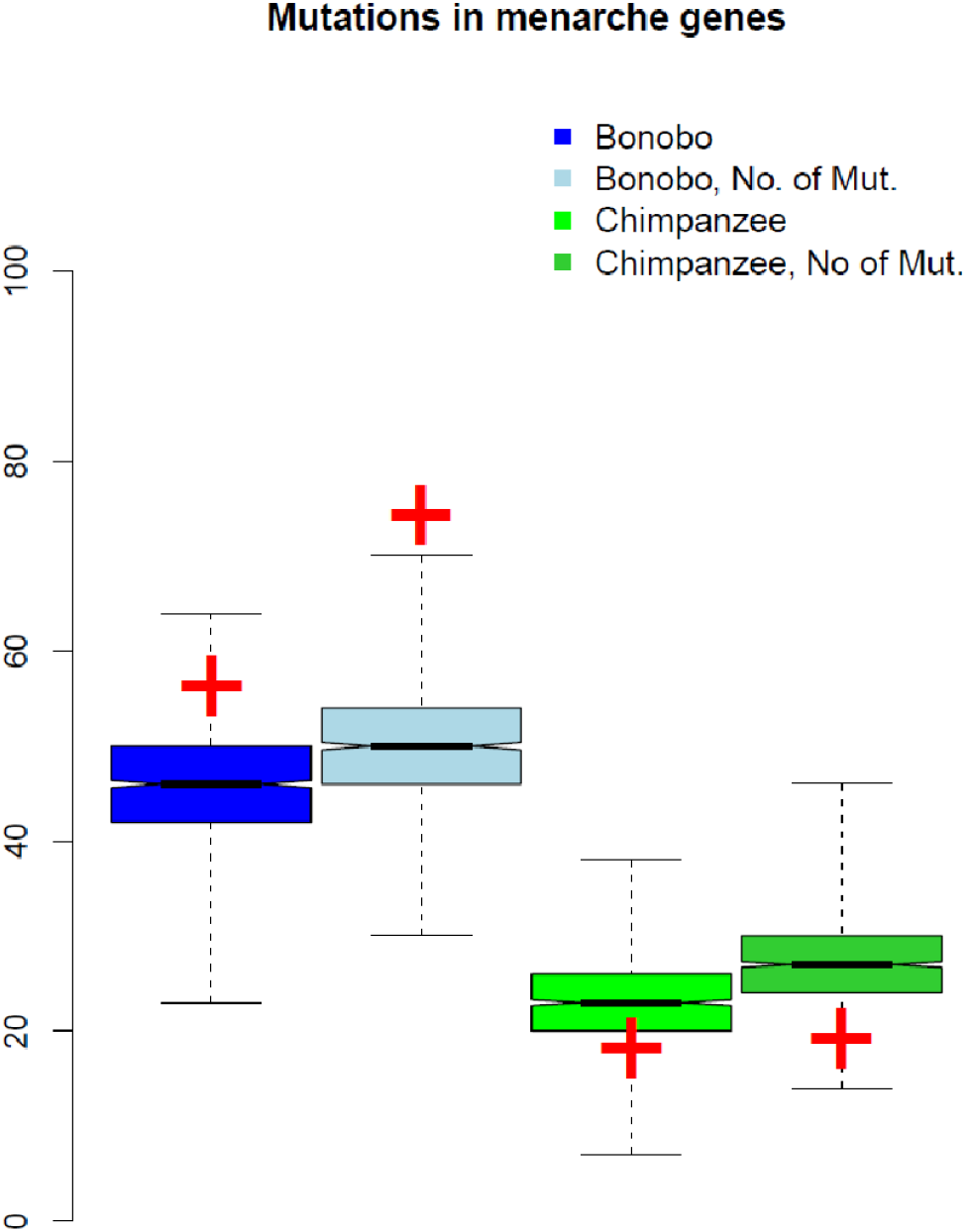
Enrichment of non-synonymous SNCs in genes associated with age at menarche in humans. The distributions of 1,000 random sets of 307 genes of similar coding length (±10%) as the menarche-associated genes are shown, with the observed values marked as red crosses. From the left, the number of genes with bonobo-specific changes, the number of bonobo-specific changes, the number of genes with chimpanzee-specific changes, the number of chimpanzee-specific changes.

## Discussion

We made use of the genetic variation across 10 bonobo and 59 chimpanzee high-coverage genomes in this study, which is the most comprehensive genomic data to date for the two *Pan* species, including all four known subspecies of chimpanzees. We use the predicted effects of single-nucleotide variants to analyze differences in the distribution of deleterious alleles between populations, stratified by protein-coding, noncoding functional and loss-of-function SNCs. We present a catalog of SNCs that are lineage-specific and determine associations to known functions in either species especially for non-synonymous variants, since these are likely to underlie phenotypic differences.

### Effective population size influences the distribution of deleterious alleles

We investigated the efficacy of purifying selection in relation to the demographic history of populations, by making use of deleterious derived alleles. Our results show that the population-wise Neutrality Index is highest in the western chimpanzee population compared to the other chimpanzee populations and bonobos (Fig. 1A). We also observe that the proportions of deleterious derived alleles, in comparison to neutral derived alleles, are highest in the western chimpanzee population across different non-coding functional element categories (Fig. 1B). The ratios of deleterious derived allele frequencies to neutral derived allele frequencies using six different deleteriousness prediction methods (Fig. 2 and Fig. S1) show that in populations which experienced a small long-term *N_e_* and population bottlenecks, proportionately more deleterious derived alleles rise and remain in the population. The proportion of deleterious derived allele frequencies in bonobos is higher at high frequencies compared to chimpanzees, which might be the consequence of a long-term small *N_e_* and multiple population bottlenecks after the split from chimpanzees (Prado-Martinez et al. 2013; De Manuel et al. 2016). At low frequencies, the proportion of deleterious derived allele frequencies is much higher in non-central chimpanzee populations compared to central chimpanzees, likely the consequence of a more recent small *N_e_* and population bottlenecks in non-central chimpanzee populations (Prado-Martinez et al. 2013; De Manuel et al. 2016). These results appear similar to the report that proportionately more deleterious derived alleles are observed in Neandertal exomes compared to modern human exomes, which diverged from each other within a similar time range as chimpanzee subpopulations did (Green et al. 2008; Briggs et al. 2009; Mendez et al. 2016), and between exomes from Eurasian and African modern humans (Castellano et al. 2014). Proportionately more deleterious derived alleles at low frequency in archaic humans and in Eurasians might be comparable to the patterns among chimpanzees, with a more recent experience of small *N_e_* and population bottlenecks, allowing deleterious mutations to rise around low frequencies. These results agree with a better efficacy to remove deleterious alleles in larger populations, as observed previously across great apes (Cagan et al. 2016), and with previous reports for modern humans, dogs and rice (Lohmueller 2014; Marsden et al. 2016; Liu et al. 2017).

On the other hand, we compare the number of homozygous to heterozygous deleterious derived alleles in each individual across populations. This clearly shows the population-wise separation in the distribution of individual deleterious load (Fig. 3A). Heterozygous sites are strongly influenced by rare variants, and a proxy for the population diversity. Homozygous derived sites, on the other hand, are likely influenced by long-term *N_e_* and population bottlenecks, and dominated by fixed or high frequency derived alleles. Although there is an ongoing discussion on this matter (Charlesworth 2013; Lohmueller 2014; Simons et al. 2014; Kim and Lohmueller 2015), population size and genetic drift seem to influence the distribution of changes in homozygous positions. This is interpreted as the influence of small *N_e_* and population bottlenecks, which causes disproportionate shifts of deleterious derived changes to high frequencies in a population (Lohmueller 2014; Simons et al. 2014). Our results on heterozygous sites suggest that the central chimpanzee population carries the largest genetic diversity, and on the homozygous sites that the western chimpanzees carry the highest level of homozygous mutational load, which appear quite similar to bonobos. We conclude that the population history generally affects the way derived changes are distributed across homo- and heterozygous sites in the genome in our closest living relatives. Interestingly, among chimpanzees, which are generally understood as a species with a large genetic diversity (Prado-Martinez et al. 2013), the western chimpanzee population appears to be rather similar to bonobos in their mutational load.

Previously, (de Valles-Ibáñez et al. 2016) analyzed the effect of population size on the deleterious burden based on LoF mutations in the great apes, suggesting it to be very weak. Here, we improve the inferences with a fine-scale analysis of the *Pan* clade and by increasing the number of individuals, which is critical for using the same number of individuals in each population when we stratify by frequency. It is important that we could include the western and central chimpanzee populations, which had the smallest and largest historical population sizes among the chimpanzee populations, respectively. The deleterious load in LoF mutations in our data agrees well with the patterns in other categories of deleterious alleles. Particularly western chimpanzees carry more fixed and homozygous LoF alleles and a relative excess of LoF over neutral singletons, which might be due to a reduced efficacy of purifying selection due to their lower population size. These observations generally support the effect of a low *N_e_* in western chimpanzees and bonobos, while the higher *N_e_* in central chimpanzees leads to a larger number of disruptive mutations in singletons, which are subsequently removed from the population by purifying selection. We conclude that in *Pan* species the effective population size does indeed have an effect on purifying selection in LoF mutations, in agreement with the observations above and in (Cagan et al. 2016).

Based on these lines of evidence, from the deleterious allele analysis in functional elements to the proportion of deleterious changes across frequencies to mutational load and LoF mutations, we conclude that the small effective population size in the past is a good proxy of a reduced efficacy of purifying selection in the *Pan* clade.

### The genetic basis of lineage-specific phenotypes

Across all deleterious categories and all loci, bonobos carry a substantially larger number of lineage-specific SNCs. This reflects that in bonobos, which experienced a long-term small *N_e_* in the past, due to genetic drift more alleles reached high frequencies or fixation. We assume that deleterious and protein-altering SNCs are likely to harbor functional consequences, possibly resulting in phenotypic changes. The concept of deleteriousness of an allele often represents conservation across species, since most new mutations are deleterious and thus removed from the population. Yet, novel functional changes would have arisen from such mutations. We present a catalog of genes with such mutations, and suggest that several of these in bonobos have functions in male and female reproduction from a literature search, and multiple genes with functions like immunity or the nervous system (Table 2).

The presence of putatively functional lineage-specific SNCs is not an immediate evidence for adaptive evolution. However, we note that several genes with non-synonymous lineage-specific SNCs (Table S7, S8) have been reported to show signatures of positive selection in bonobos (Cagan et al. 2016): *PCDH15, IQCA1, CCDC149* and *SLC36A1* in the Fay and Wu’s H test, and *CFH* and *ULK4* in the HKA based test (Table S9). *PCDH15* (with two non-synonymous changes in bonobos, one of them predicted to be deleterious by SIFT and PolyPhen-2) is involved in retinal and cochlear function (Jacobson et al. 2008), and *ULK4* has been associated with schizophrenia and neuronal function (Lang et al. 2014). Among the genes with non-synonymous lineage-specific SNCs in chimpanzees *MUC13* and *ADGB* were described to show signatures of positive selection (ELS test; Cagan et al. 2016).

One of the most prominent differences between chimpanzees and bonobos is found in female reproduction, with female bonobos having prolonged maximal sexual swellings, which female chimpanzees do not have, and female bonobos experiencing an earlier onset of their reproductive age. We hypothesized that SNCs on the bonobo lineage influencing the female reproductive system might underlie the phenotypic differences of the two species. Genome-wide signatures of strong positive selection (Cagan et al. 2016) found no enrichment in female reproduction genes, which could be due to the limited power of these methods to detect selection early after the divergence of the two species. However, we demonstrate that the trait “age at menarche” is among only five traits significantly enriched for bonobo-specific non-synonymous changes. Furthermore, the most complete dataset of genes associated to this trait shows an even stronger enrichment in bonobos, with 73 protein-coding SNCs in 55 genes (Table 2). Most of these genes (98%) are expressed in primary tissues of chimpanzees and bonobos (Table S13), suggesting that they are functional in the *Pan* clade. Among the 307 menarche-related genes, five (*ATE1, DLGAP1, CSMD1, LRP1B, TRPC6*) have been reported to show signatures of positive selection (Cagan et al. 2016), and one (*HLA-DQB1*) of balancing selection in bonobos. Among these, the Low Density Lipoprotein Receptor *LRP1B* carries three protein-coding changes, and *CSMD1* carries one (Table S13). We additionally find one nonsense SNC in *RPRD2* and one start codon SNC in *NKAIN2*. These genes constitute the strongest single candidates for lineage-specific functional changes.

To our knowledge, this is the first time that this complex trait in female bonobos has been investigated using genetic data. The main limitation of this study is that only ten bonobo genomes are available. Bonobos are an understudied species in population genetics, hence fine-scaled studies of their population structure and sequencing of more individuals would improve power in future studies. Despite age at menarche being the associated trait in humans, these genes encompass broader functions in the female reproductive system, rather than controlling only age at menarche. Hence, we interpret this as an enrichment of functional changes in female reproduction-related genes during the evolutionary history of bonobos on a polygenic scale, with *LRP1B* and *CSMD1* being good candidates to have the strongest influence. Our results are in agreement with suggestions that the prominent physiological differences between chimpanzee and bonobo female sexual swellings could be due to derived features in bonobos (Wrangham 1993), and suggest that it might have been adaptive on the bonobo lineage. These bonobo-specific non-synonymous changes in menarche-related genes deserve further investigation on the functional level, which would serve as the foundation for a better understanding of the female reproductive system. The sexual swelling in female bonobos has a profound influence on their biology and group dynamics (Hohmann and Fruth 2000; Furuichi 2011), which can be considered typical for bonobo-specific behavior and sociality. Since other relevant traits show an enrichment of changes on the bonobo lineage as well (*e.g*. in behavior- and cognition-related genes), these deserve further investigation in future studies.

## Acknowledgements

We thank Marc de Manuel for help with data preparation and technical advice. This work was supported by the Max Planck Society [to AMA], BFU2017-86471-P (MINECO/FEDER, UE) [to TMB], U01 [MH106874 to TMB], Howard Hughes International Early Career [to TMB], Obra Social “La Caixa” [to TMB], Secretaria d’Universitats i Recerca del Departament d’Economia i Coneixement de la Generalitat de Catalunya [to TMB], and a Deutsche Forschungsgemeinschaft (DFG) fellowship [KU 3467/1-1 to MK].

## Author Contributions

TMB, AMA and MK conceived the project and wrote the paper. SH and MK analyzed data and wrote the paper.

## References

Adzhubei IA, Schmidt S, Peshkin L, Ramensky VE, Gerasimova A, Bork P, Kondrashov AS, Sunyaev SR. 2010. A method and server for predicting damaging missense mutations. Nat. Methods [Internet] 7:248–249. Available from: http://www.ncbi.nlm.nih.gov/pubmed/20354512

Anders S, Pyl PT, Huber W. 2015. HTSeq—a Python framework to work with high-throughput sequencing data. Bioinformatics 31:166.

Bataillon T, Duan J, Hvilsom C, Jin X, Li Y, Skov L, Glemin S, Munch K, Jiang T, Qian Y, et al. 2015. Inference of Purifying and Positive Selection in Three Subspecies of Chimpanzees (Pan troglodytes) from Exome Sequencing. Genome Biol. Evol. 7:1122–1132.

Behringer V, Deschner T, Deimel C, Stevens JMG, Hohmann G. 2014. Age-related changes in urinary testosterone levels suggest differences in puberty onset and divergent life history strategies in bonobos and chimpanzees. Horm. Behav. [Internet] 66:525–533. Available from: http://www.ncbi.nlm.nih.gov/pubmed/25086337

Brawand D, Soumillon M, Necsulea A, Julien P, Csárdi G, Harrigan P, Weier M, Liechti A, Aximu-Petri A, Kircher M, et al. 2011. The evolution of gene expression levels in mammalian organs. [SupMat]. Nature 478:343–348.

Briggs AW, Good JM, Green RE, Krause J, Maricic T, Stenzel U, Lalueza-Fox C, Rudan P, Brajkovic D, Kucan Z, et al. 2009. Targeted Retrieval and Analysis of Five Neandertal mtDNA Genomes. Science (80-.). [Internet] 325:318–321. Available from: http://www.ncbi.nlm.nih.gov/pubmed/19608918

Cagan A, Theunert C, Laayouni H, Santpere G, Pybus M, Casals F, Prüfer K, Navarro A, Marques-Bonet T, Bertranpetit J, et al. 2016. Natural Selection in the Great Apes. Mol. Biol. Evol. [Internet] 33:3268–3283. Available from: https://academic.oup.com/mbe/article-lookup/doi/10.1093/molbev/msw215

Castellano S, Parra G, Sánchez-Quinto FA, Racimo F, Kuhlwilm M, Kircher M, Sawyer S, Fu Q, Heinze A, Nickel B, et al. 2014. Patterns of coding variation in the complete exomes of three Neandertals.

Charlesworth B. 2013. WHY WE ARE NOT DEAD ONE HUNDRED TIMES OVER. Evolution (N. Y). 67:3354–3361.

Daisuke A, Uoshinao O, Satoshi H, Ken-ichi T, Yoshihiro O, Yuji B, Shinji O, Nao S, Suminori K, Masazumi T, et al. 2009. Overexpression of class III beta-tubulin predicts good response to taxane-based chemotherapy in ovarian clear cell adenocarcinoma. Clin. Cancer Res. [Internet] 15:1473–1480. Available from: http://www.ncbi.nlm.nih.gov/pubmed/19228748

Davydov E V., Goode DL, Sirota M, Cooper GM, Sidow A, Batzoglou S. 2010. Identifying a High Fraction of the Human Genome to be under Selective Constraint Using GERP++.Wasserman WW, editor. PLoS Comput. Biol. [Internet] 6:e1001025. Available from: http://dx.plos.org/10.1371/journal.pcbi.1001025

Day FR, Thompson DJ, Helgason H, Chasman DI, Finucane H, Sulem P, Ruth KS, Whalen S, Sarkar AK, Albrecht E, et al. 2017. Genomic analyses identify hundreds of variants associated with age at menarche and support a role for puberty timing in cancer risk. Nat. Genet. 49:834–841.

Durinck S, Moreau Y, Kasprzyk A, Davis S, De Moor B, Brazma A, Huber W. 2005. BioMart and Bioconductor: a powerful link between biological databases and microarray data analysis. Bioinformatics 21:3439–3440.

Eyre-Walker A. 2006. The genomic rate of adaptive evolution. Trends Ecol. Evol. 21:569–575.

Fischer A, Prüfer K, Good JM, Halbwax M, Wiebe V, André C, Atencia R, Mugisha L, Ptak SE, Pääbo S. 2011. Bonobos Fall within the Genomic Variation of Chimpanzees.Joly E, editor. PLoS One [Internet] 6:e21605. Available from: http://dx.plos.org/10.1371/journal.pone.0021605

Fraisl P, Tanaka H, Forss-Petter S, Lassmann H, Nishimune Y, Berger J. 2006. A novel mammalian bubblegum-related acyl-CoA synthetase restricted to testes and possibly involved in spermatogenesis. Arch. Biochem. Biophys. [Internet] 451:23–33. Available from: http://linkinghub.elsevier.com/retrieve/pii/S0003986106001615

Furuichi T. 1987. Sexual swelling, receptivity, and grouping of wild pygmy chimpanzee females at Wamba, Zaïre. Primates [Internet] 28:309–318. Available from: http://link.springer.com/10.1007/BF02381014

Furuichi T. 2011. Female contributions to the peaceful nature of bonobo society. Evol. Anthropol. Issues, News, Rev. [Internet] 20:131–142. Available from: http://doi.wiley.com/10.1002/evan.20308

Goodall J. 1986. The Chimpanzees of Gombe: Patterns of Behavior. Press Harvard Univ. [Internet]. Available from: https://repository.library.georgetown.edu/handle/10822/811357

Grantham R. 1974. Amino Acid Difference Formula to Help Explain Protein Evolution. Science (80-.). 185.

Green RE, Malaspinas A-S, Krause J, Briggs AW, Johnson PLF, Uhler C, Meyer M, Good JM, Maricic T, Stenzel U, et al. 2008. A Complete Neandertal Mitochondrial Genome Sequence Determined by High-Throughput Sequencing. Cell [Internet] 134:416–426. Available from: http://www.ncbi.nlm.nih.gov/pubmed/18692465

Gronau I, Hubisz MJ, Gulko B, Danko CG, Siepel A. 2011. Bayesian inference of ancient human demography from individual genome sequences. Nat. Genet. 43:1031–1034.

Gurvits N, Löyttyniemi E, Nykänen M, Kuopio T, Kronqvist P, Talvinen K. 2017. Separase is a marker for prognosis and mitotic activity in breast cancer. Br. J. Cancer [Internet] 117:1383–1391. Available from: http://www.nature.com/doifinder/10.1038/bjc.2017.301

Hare B, Wobber V, Wrangham R. 2012. The self-domestication hypothesis: evolution of bonobo psychology is due to selection against aggression. Anim. Behav. [Internet] 83:573–585. Available from: http://linkinghub.elsevier.com/retrieve/pii/S000334721100546X

Henn BM, Botigué LR, Peischl S, Dupanloup I, Lipatov M, Maples BK, Martin AR, Musharoff S, Cann H, Snyder MP, et al. 2016. Distance from sub-Saharan Africa predicts mutational load in diverse human genomes. Proc. Natl. Acad. Sci. U. S. A. [Internet] 113:E440–9. Available from: http://www.ncbi.nlm.nih.gov/pubmed/26712023

Hering D, Olenski K, Kaminski S. 2014. Genome-Wide Association Study for Sperm Concentration in Holstein-Friesian Bulls. Reprod. Domest. Anim. [Internet] 49:1008–1014. Available from: http://doi.wiley.com/10.1111/rda.12423

Hodgkinson A, Casals F, Idaghdour Y, Grenier J-C, Hernandez RD, Awadalla P. 2013. Selective constraint, background selection, and mutation accumulation variability within and between human populations. BMC Genomics [Internet] 14:495. Available from: http://www.ncbi.nlm.nih.gov/pubmed/23875710

Hohmann G, Fruth B. 2000. Use and function of genital contacts among female bonobos. Anim. Behav. [Internet] 60:107–120. Available from: http://www.ncbi.nlm.nih.gov/pubmed/10924210

Hozumi A, Padma P, Toda T, Ide H, Inaba K. 2008. Molecular characterization of axonemal proteins and signaling molecules responsible for chemoattractant-induced sperm activation inCiona intestinalis. Cell Motil. Cytoskeleton [Internet] 65:249–267. Available from: http://doi.wiley.com/10.1002/cm.20258

Huber W, Carey VJ, Gentleman R, Anders S, Carlson M, Carvalho BS, Bravo HC, Davis S, Gatto L, Girke T, et al. 2015. Orchestrating high-throughput genomic analysis with Bioconductor. Nat Meth 12:115–121.

Idani G. 1990. RELATIONS BETWEEN UNIT-GROUPS OF BONOBOS AT WAMBA, ZAIRE: ENCOUNTERS AND TEMPORARY FUSIONS. Afr. Study Monogr. [Internet]. Available from: http://www.africa.kyoto-u.ac.jp/kiroku/asm_normal/abstracts/pdf/ASM Vol.11 No.3 1990/WWW Genichi IDANI.pdf

Jacobson SG, Cideciyan A V, Aleman TS, Sumaroka A, Roman AJ, Gardner LM, Prosser HM, Mishra M, Bech-Hansen NT, Herrera W, et al. 2008. Usher syndromes due to MYO7A, PCDH15, USH2A or GPR98 mutations share retinal disease mechanism. Hum. Mol. Genet. 17:2405–2415.

Jukes TH, Kimura M. 1984. Evolutionary constraints and the neutral theory. J. Mol. Evol. [Internet] 21:90–92. Available from: http://link.springer.com/10.1007/BF02100633

Kidd JM, Gravel S, Byrnes J, Moreno-Estrada A, Musharoff S, Bryc K, Degenhardt JD, Brisbin A, Sheth V, Chen R, et al. 2012. Population genetic inference from personal genome data: impact of ancestry and admixture on human genomic variation. Am. J. Hum. Genet. [Internet] 91:660–671. Available from: http://www.ncbi.nlm.nih.gov/pubmed/23040495

Kim BY, Lohmueller KE. 2015. Selection and reduced population size cannot explain higher amounts of Neandertal ancestry in East Asian than in European human populations. Am. J. Hum. Genet. 96:454–461.

Kim D, Pertea G, Trapnell C, Pimentel H, Kelley R, Salzberg SL. 2013. TopHat2: accurate alignment of transcriptomes in the presence of insertions, deletions and gene fusions. Genome Biol. 14:R36.

Kircher M, Witten DM, Jain P, O’Roak BJ, Cooper GM, Shendure J. 2014. A general framework for estimating the relative pathogenicity of human genetic variants. Nat. Genet. 46:310–315.

Konecny GE, Pauletti G, Untch M, Wang H-J, Möbus V, Kuhn W, Thomssen C, Harbeck N, Wang L, Apple S, et al. 2010. Association between HER2, TOP2A, and response to anthracycline-based preoperative chemotherapy in high-risk primary breast cancer. Breast Cancer Res. Treat. [Internet] 120:481–489. Available from: http://link.springer.com/10.1007/s10549-010-0744-z

Kuhlwilm M, Manuel M de, Nater A, Greminger MP, Krützen M, Marques-Bonet T. 2016. Evolution and demography of the great apes. Curr. Opin. Genet. Dev.

Kumar P, Henikoff S, Ng PC. 2009. Predicting the effects of coding non-synonymous variants on protein function using the SIFT algorithm. Nat. Protoc. [Internet] 4:1073–1081. Available from: http://www.ncbi.nlm.nih.gov/pubmed/19561590

Kuroda S. 1980. Social behavior of the pygmy chimpanzees. Primates [Internet] 21:181–197. Available from: http://link.springer.com/10.1007/BF02374032

Lang B, Pu J, Hunter I, Liu M, Martin-Granados C, Reilly TJ, Gao G-D, Guan Z-L, Li W-D, Shi Y-Y, et al. 2014. Recurrent deletions of ULK4 in schizophrenia: a gene crucial for neuritogenesis and neuronal motility. J. Cell Sci. 127:630–640.

Langergraber K, Prüfer K. 2012. Generation times in wild chimpanzees and gorillas suggest earlier divergence times in great ape and human evolution. Proc.…109:15716–15721.

Lawrence M, Huber W, Pagès H, Aboyoun P, Carlson M, Gentleman R, Morgan MT, Carey VJ. 2013. Software for Computing and Annotating Genomic Ranges. PLOS Comput. Biol. 9:e1003118.

Lek M, Karczewski KJ, Minikel E V, Samocha KE, Banks E, Fennell T, O’Donnell-Luria AH, Ware JS, Hill AJ, Cummings BB, et al. 2016. Analysis of protein-coding genetic variation in 60,706 humans. Nature 536:285–291.

Li H, Handsaker B, Wysoker A, Fennell T, Ruan J, Homer N, Marth G, Abecasis G, Durbin R. 2009. The Sequence Alignment/Map format and SAMtools. Bioinformatics 25:2078–2079.

Li W-H, Wu C-I, Luo C-C. 1984. Nonrandomness of point mutation as reflected in nucleotide substitutions in pseudogenes and its evolutionary implications.J. Mol. Evol. [Internet] 21:58–71. Available from: http://link.springer.com/10.1007/BF02100628

Liu Q, Zhou Y, Morrell PL, Gaut BS. 2017. Deleterious variants in Asian rice and the potential cost of domestication. Mol. Biol. Evol. [Internet] 167:msw296. Available from: https://academic.oup.com/mbe/article-lookup/doi/10.1093/molbev/msw296

Lohmueller KE. 2014. The distribution of deleterious genetic variation in human populations. Curr. Opin. Genet. Dev. [Internet] 29:139–146. Available from: http://linkinghub.elsevier.com/retrieve/pii/S0959437X14001002

Lohmueller KE, Indap AR, Schmidt S, Boyko AR, Hernandez RD, Hubisz MJ, Sninsky JJ, White TJ, Sunyaev SR, Nielsen R, et al. 2008. Proportionally more deleterious genetic variation in European than in African populations. Nature [Internet] 451:994–997. Available from: http://www.nature.com/doifinder/10.1038/nature06611

MacArthur J, Bowler E, Cerezo M, Gil L, Hall P, Hastings E, Junkins H, McMahon A, Milano A, Morales J, et al. 2017. The new NHGRI-EBI Catalog of published genome-wide association studies (GWAS Catalog). Nucleic Acids Res. [Internet] 45:D896–D901. Available from: http://www.ncbi.nlm.nih.gov/pubmed/27899670

De Manuel M, Kuhlwilm M, Frandsen P, Sousa VC, Desai T, Prado-Martinez J, Hernandez-Rodriguez J, Dupanloup I, Lao O, Hallast P, et al. 2016. Chimpanzee genomic diversity reveals ancient admixture with bonobos. Science (80-.). 354.

Marsden CD, Ortega-Del Vecchyo D, O’Brien DP, Taylor JF, Ramirez O, Vilà C, Marques-Bonet T, Schnabel RD, Wayne RK, Lohmueller KE. 2016. Bottlenecks and selective sweeps during domestication have increased deleterious genetic variation in dogs. Proc. Natl. Acad. Sci. U. S. A. [Internet] 113:152–157. Available from: http://www.ncbi.nlm.nih.gov/pubmed/26699508

McLaren W, Gil L, Hunt SE, Riat HS, Ritchie GRS, Thormann A, Flicek P, Cunningham F. 2016. The Ensembl Variant Effect Predictor. Genome Biol. 17:122.

Mendez FL, Poznik GD, Castellano S, Bustamante CD. 2016. The Divergence of Neandertal and Modern Human Y Chromosomes. Am. J. Hum. Genet. [Internet] 98:728–734. Available from: http://www.ncbi.nlm.nih.gov/pubmed/27058445

Nishida T. 1983. Alpha status and agonistic alliance in wild chimpanzees (Pan troglodytes schweinfurthii). Primates [Internet] 24:318–336. Available from: http://link.springer.com/10.1007/BF02381978

Nunn CL. 1999. The evolution of exaggerated sexual swellings in primates and the graded-signal hypothesis. Anim. Behav. [Internet] 58:229–246. Available from: http://www.ncbi.nlm.nih.gov/pubmed/10458874

Nymoen DA, Holth A, Hetland Falkenthal TE, Tropé CG, Davidson B. 2015. CIAPIN1 and ABCA13 are markers of poor survival in metastatic ovarian serous carcinoma. Mol. Cancer [Internet] 14:44. Available from: http://www.ncbi.nlm.nih.gov/pubmed/25889687

Nystad M, Sitras V, Larsen M, Acharya G. 2014. Placental expression of aminopeptidase-Q (laeverin) and its role in the pathophysiology of preeclampsia. Am. J. Obstet. Gynecol. [Internet] 211:686.e1-686.e31. Available from: http://www.ncbi.nlm.nih.gov/pubmed/24959655

Prado-Martinez J, Sudmant PH, Kidd JM, Li H, Kelley JL, Lorente-Galdos B, Veeramah KR, Woerner AE, O’Connor TD, Santpere G, et al. 2013. Great ape genetic diversity and population history. Nature [Internet] 499:471–475. Available from: http://www.ncbi.nlm.nih.gov/pubmed/23823723

Prüfer K, Muetzel B, Do H-H, Weiss G, Khaitovich P, Rahm E, Pääbo S, Lachmann M, Enard W. 2007. FUNC: a package for detecting significant associations between gene sets and ontological annotations. BMC Bioinformatics 8:41.

Rand DM, Kann LM. 1996. Excess amino acid polymorphism in mitochondrial DNA: contrasts among genes from Drosophila, mice, and humans. Mol. Biol. Evol. 13:735–748.

Randall Parish A. 1994. Sex and food control in the “uncommon chimpanzee”: How Bonobo females overcome a phylogenetic legacy of male dominance. Ethol. Sociobiol. [Internet] 15:157–179. Available from: http://www.sciencedirect.com/science/article/pii/0162309594900388

Ritchie GRS, Dunham I, Zeggini E, Flicek P. 2014. Functional annotation of noncoding sequence variants. Nat. Methods [Internet] 11:294–296. Available from: http://www.nature.com/doifinder/10.1038/nmeth.2832

Ryu H, Hill DA, Furuichi T. 2015. Prolonged maximal sexual swelling in wild bonobos facilitates affiliative interactions between females. Behaviour [Internet] 152:285–311. Available from: http://booksandjournals.brillonline.com/content/journals/10.1163/1568539x-00003212

Sadek CM, Damdimopoulos AE, Pelto-Huikko M, Gustafsson J-A, Spyrou G, Miranda-Vizuete A. 2001. Sptrx-2, a fusion protein composed of one thioredoxin and three tandemly repeated NDP-kinase domains is expressed in human testis germ cells. Genes to Cells [Internet] 6:1077–1090. Available from: http://doi.wiley.com/10.1046/j.1365-2443.2001.00484.x

Scanlan MJ, Gout I, Gordon CM, Williamson B, Stockert E, Gure AO, Jäger D, Chen YT, Mackay A, O’Hare MJ, et al. 2001. Humoral immunity to human breast cancer: antigen definition and quantitative analysis of mRNA expression. Cancer Immun. 1:4.

Simons YB, Turchin MC, Pritchard JK, Sella G. 2014. The deleterious mutation load is insensitive to recent population history.Nat. Genet. 46:220–224.

Singel SM, Cornelius C, Zaganjor E, Batten K, Sarode VR, Buckley DL, Peng Y, John GB, Li HC, Sadeghi N, et al. 2014. KIF14 Promotes AKT Phosphorylation and Contributes to Chemoresistance in Triple-Negative Breast Cancer. Neoplasia [Internet] 16:247–256.e2. Available from: http://www.ncbi.nlm.nih.gov/pubmed/24784001

Subramanian S, Lambert DM. 2012. Selective constraints determine the time dependency of molecular rates for human nuclear genomes. Genome Biol. Evol. [Internet] 4:1127–1132. Available from: http://www.ncbi.nlm.nih.gov/pubmed/23059453

Takasaki N, Tachibana K, Ogasawara S, Matsuzaki H, Hagiuda J, Ishikawa H, Mochida K, Inoue K, Ogonuki N, Ogura A, et al. 2014. A heterozygous mutation of GALNTL5 affects male infertility with impairment of sperm motility. Proc. Natl. Acad. Sci. U. S. A. [Internet] 111:1120–1125. Available from: http://www.ncbi.nlm.nih.gov/pubmed/24398516

Takemoto H, Kawamoto Y, Higuchi S, Makinose E, Hart JA, Hart TB, Sakamaki T, Tokuyama N, Reinartz GE, Guislain P, et al. 2017. The mitochondrial ancestor of bonobos and the origin of their major haplogroups. PLoS One [Internet] 12:e0174851. Available from: http://www.ncbi.nlm.nih.gov/pubmed/28467422

Thompson-Handler N, Malenky RK, Badrian N. 1984. Sexual Behavior of Pan paniscus under Natural Conditions in the Lomako Forest, Equateur, Zaire. In: The Pygmy Chimpanzee. Boston, MA: Springer US. p. 347–368. Available from: http://link.springer.com/10.1007/978-1-4757-0082-4_14

Thompson JAM. 2003. A model of the biogeographical journey from Proto-pan to Pan paniscus. Primates [Internet] 44:191–197. Available from: https://link.springer.com/article/10.1007/s10329-002-0029-1

Torkamani A, Pham P, Libiger O, Bansal V, Zhang G, Scott-Van Zeeland AA, Tewhey R, Topol EJ, Schork NJ. 2012. Clinical implications of human population differences in genome-wide rates of functional genotypes. Front. Genet. [Internet] 3:211. Available from: http://www.ncbi.nlm.nih.gov/pubmed/23125845

Tubbs R, Barlow WE, Budd GT, Swain E, Porter P, Gown A, Yeh I-T, Sledge G, Shapiro C, Ingle J, et al. 2009. Outcome of patients with early-stage breast cancer treated with doxorubicin-based adjuvant chemotherapy as a function of HER2 and TOP2A status. J. Clin. Oncol. [Internet] 27:3881–3886. Available from: http://www.ncbi.nlm.nih.gov/pubmed/19620488

de Valles-Ibáñez G, Hernandez-Rodriguez J, Prado-Martinez J, Luisi P, Marquès-Bonet T, Casals F. 2016. Genetic load of loss-of-function polymorphic variants in great apes. Genome Biol. Evol.:evw040.

Wong JCY. 2003. Targeted disruption of exons 1 to 6 of the Fanconi Anemia group A gene leads to growth retardation, strain-specific microphthalmia, meiotic defects and primordial germ cell hypoplasia. Hum. Mol. Genet. [Internet] 12:2063–2076. Available from: https://academic.oup.com/hmg/article-lookup/doi/10.1093/hmg/ddg219

Wrangham RW. 1993. The evolution of sexuality in chimpanzees and bonobos. Hum. Nat. [Internet] 4:47–79. Available from: http://link.springer.com/10.1007/BF02734089

Wu ATH, Sutovsky P, Manandhar G, Xu W, Katayama M, Day BN, Park K-W, Yi Y-J, Xi YW, Prather RS, et al. 2007. PAWP, a Sperm-specific WW Domain-binding Protein, Promotes Meiotic Resumption and Pronuclear Development during Fertilization. J. Biol. Chem. [Internet] 282:12164–12175. Available from: http://www.jbc.org/lookup/doi/10.1074/jbc.M609132200

Xue Y, Prado-Martinez J, Sudmant PH, Narasimhan V, Ayub Q, Szpak M, Frandsen P, Chen Y, Yngvadottir B, Cooper DN, et al. 2015. Mountain gorilla genomes reveal the impact of long-term population decline and inbreeding. Science [Internet] 348:242–245. Available from: http://www.ncbi.nlm.nih.gov/pubmed/25859046

